## Supplementary Data for "Structural basis of gap-filling DNA synthesis in the nucleosome by DNA Polymerase β"

### Supplementary Fig. 1

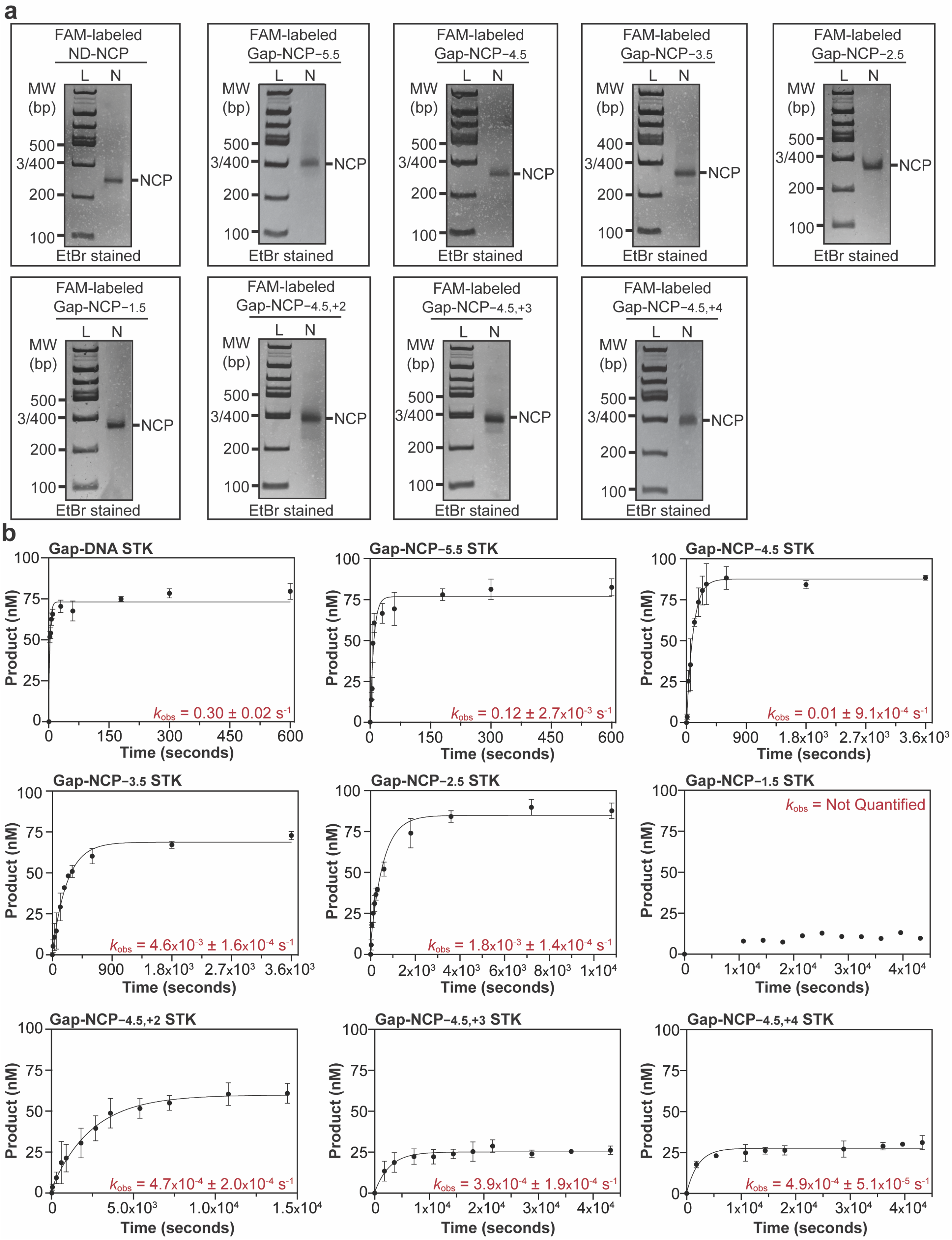

**Supplementary Fig. 1: Analysis of Pol β nucleotide insertion rate for Gap-NCPs**

**a,** Native PAGE gels confirming nucleosome formation and purity for the 6-FAM-labeled ND-NCP, Gap-NCP$_{-5.5}$, Gap-NCP$_{-4.5}$, Gap-NCP$_{-3.5}$, Gap-NCP$_{-2.5}$, Gap-NCP$_{-1.5}$, Gap-NCP$_{-4.5,+2}$, Gap-NCP$_{-4.5,+3}$, and Gap-NCP$_{-4.5,+4}$ samples. The native PAGE gels were immediately run after initial nucleosome formation and purification (n=1). The NCPs were detected using ethidium bromide staining. The 100 bp DNA ladder (L) and nucleosome sample (N) are labeled. **b,** Quantification and fits for the Gap-DNA, Gap-NCP$_{-5.5}$, Gap-NCP$_{-4.5}$, Gap-NCP$_{-3.5}$, Gap-NCP$_{-2.5}$, Gap-NCP$_{-1.5}$, Gap-NCP$_{-4.5,+2}$, Gap-NCP$_{-4.5,+3}$, and Gap-NCP$_{-4.5,+4}$ single turnover kinetic (STK) experiments. The data points represent the mean ± standard deviation from three independent replicate experiments. The error bars are included for all experimental data points, but some error bars are smaller than the circles used to represent the data points. The kinetic parameter $k_{obs}$ is shown as an inset for each experiment and represents the mean ± standard error of the mean from the three independent replicate experiments. The kinetic parameter $k_{obs}$ was not quantified for Gap-NCP$_{-1.5}$ due to minimal product formation (~10%). All source data in this figure are provided as a Source Data file.

**Supplementary Fig. 2**

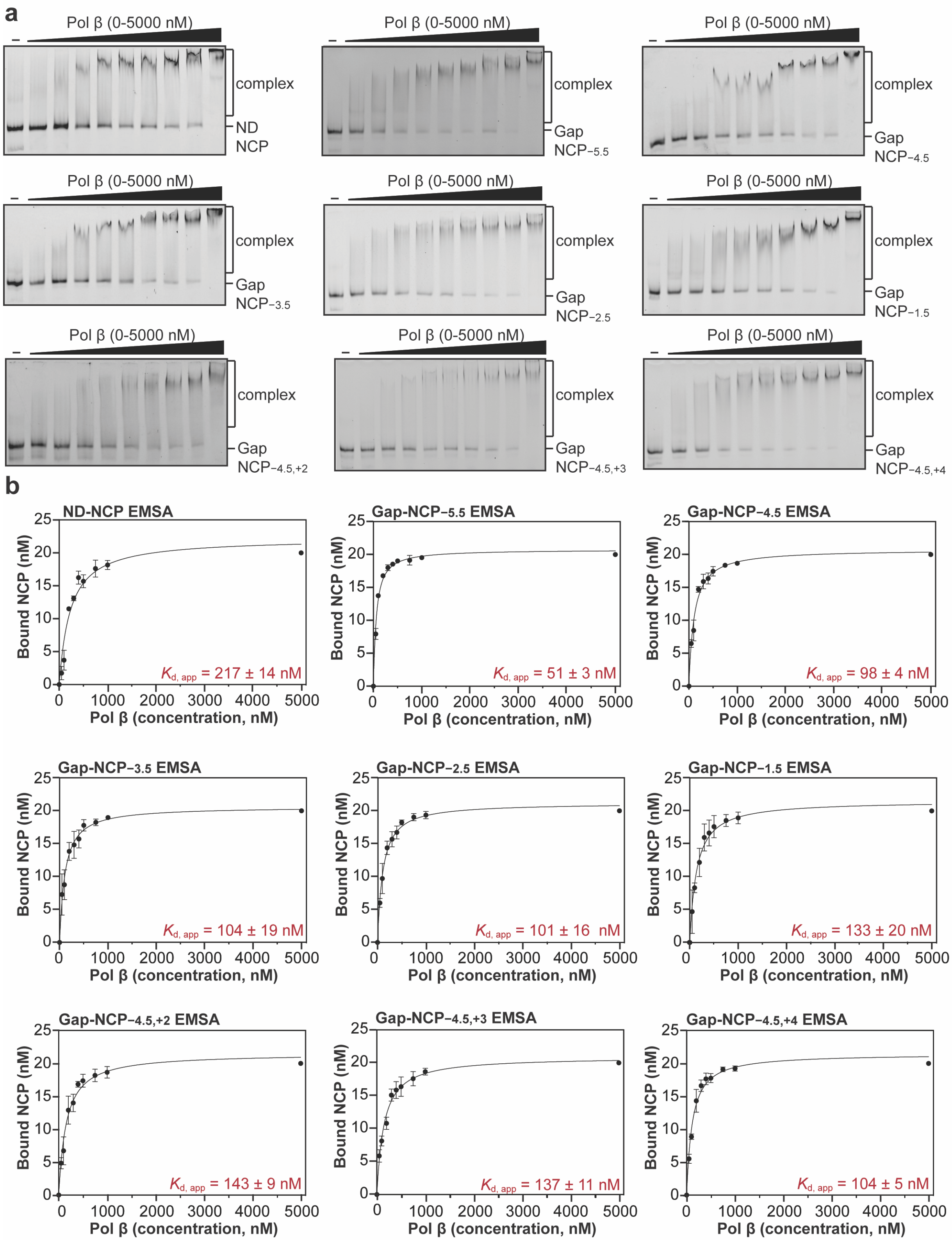

**Supplementary Fig. 2: Analysis of Pol β binding affinity for Gap-NCPs**

**a,** Representative native PAGE gels from electrophoretic mobility shift assays (EMSAs) of Pol β and ND-NCP, Gap-NCP$_{-5.5}$, Gap-NCP$_{-4.5}$, Gap-NCP$_{-3.5}$, Gap-NCP$_{-2.5}$, Gap-NCP$_{-1.5}$, Gap-NCP$_{-4.5,+2}$, Gap-NCP$_{-4.5,+3}$, and Gap-NCP$_{-4.5,+4}$. These gels are representative of three independent replicate EMSA experiments for each NCP. The free nucleosome and complex were detected using the 6-FAM label on each NCP**. b,** Quantification and fits from the EMSA experiments with Pol β and ND-NCP, Gap-NCP$_{-5.5}$, Gap-NCP$_{-4.5}$, Gap-NCP$_{-3.5}$, Gap-NCP$_{-2.5}$, Gap-NCP$_{-1.5}$, Gap-NCP$_{-4.5,+2}$, Gap-NCP$_{-4.5,+3}$, and Gap-NCP$_{-4.5,+4}$. The data points represent the mean ± standard deviation from three independent replicate experiments. The error bars are included for all experimental data points, but some error bars are smaller than the circles used to represent the data points. The apparent binding affinity ($K_{d,\ app}$) is shown as an inset for each experiment and represents the mean ± standard error of the mean from the three independent replicate experiments. All source data in this figure are provided as a Source Data file.

#### Supplementary Fig. 3

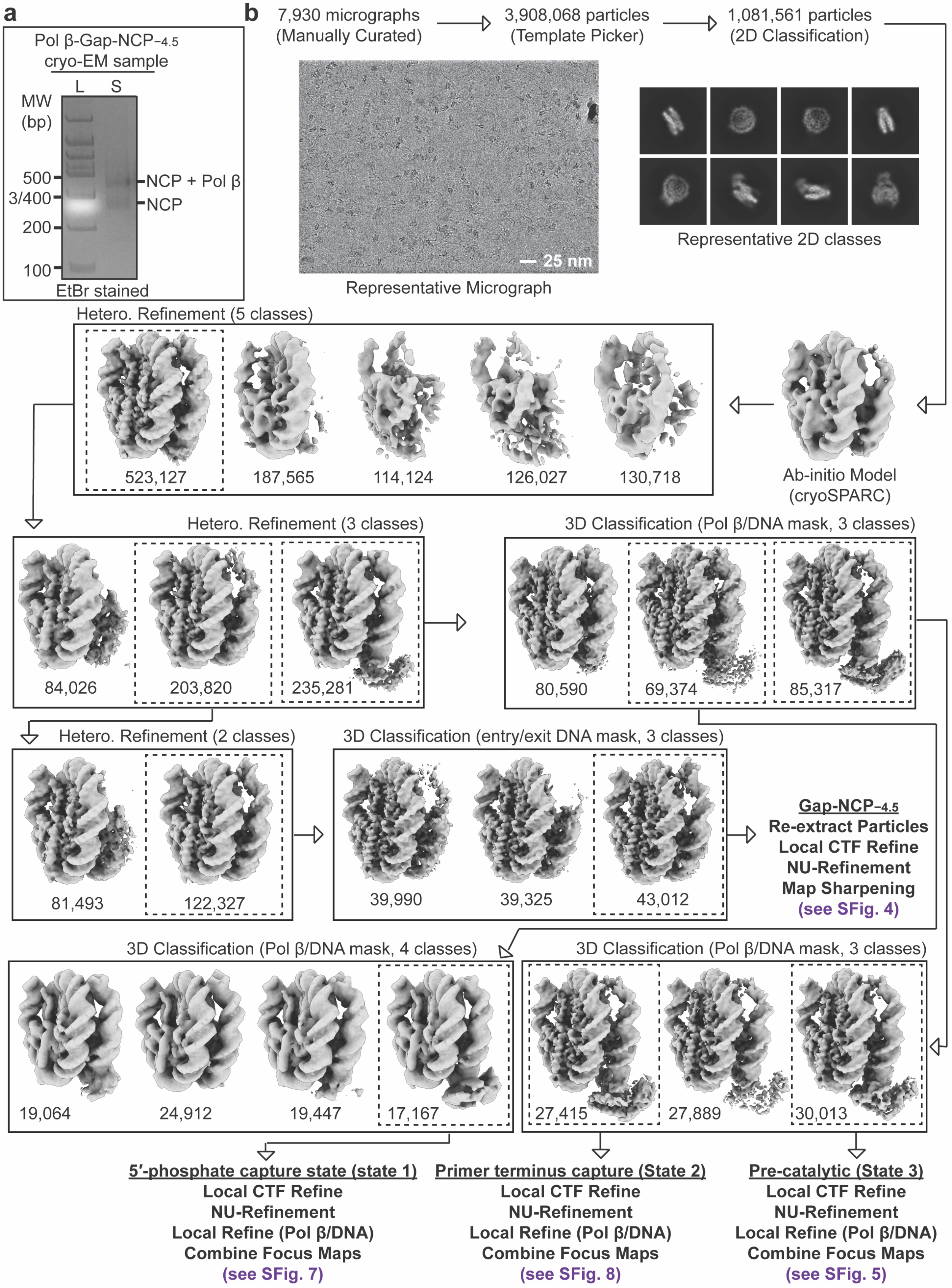

**Supplementary Fig. 3: Pol β-Gap-NCP$_{-4.5}$ SPA processing workflow**

**a,** Native PAGE gel of the Pol β-Gap-NCP$_{-4.5}$ cryo-EM sample (S) and a 100 bp DNA ladder (L). The Gap-NCP$_{-4.5}$ and Pol β-Gap-NCP$_{-4.5}$ complex were visualized with ethidium bromide staining. **b,** Flowchart of the data processing pipeline for the Pol β-Gap-NCP$_{-4.5}$ cryo-EM dataset. A representative micrograph (n=7,930) and representative 2D classes from the Pol β-Gap-NCP$_{-4.5}$ cryo-EM dataset are shown. The maps chosen for further classification and/or refinement throughout the data processing pipeline are boxed. The final maps, final models, and quality assessment metrics for Gap-NCP$_{-4.5}$, Pol β-Gap-NCP$_{-4.5}$ (5′-phosphate capture, state 1), Pol β-Gap-NCP$_{-4.5}$ (primer terminus capture, state 2), and Pol β-Gap-NCP$_{-4.5}$ (pre-catalytic, state 3) can be found in Supplementary Figs. 4,7,8,5, respectively.

#### Supplementary Fig. 4

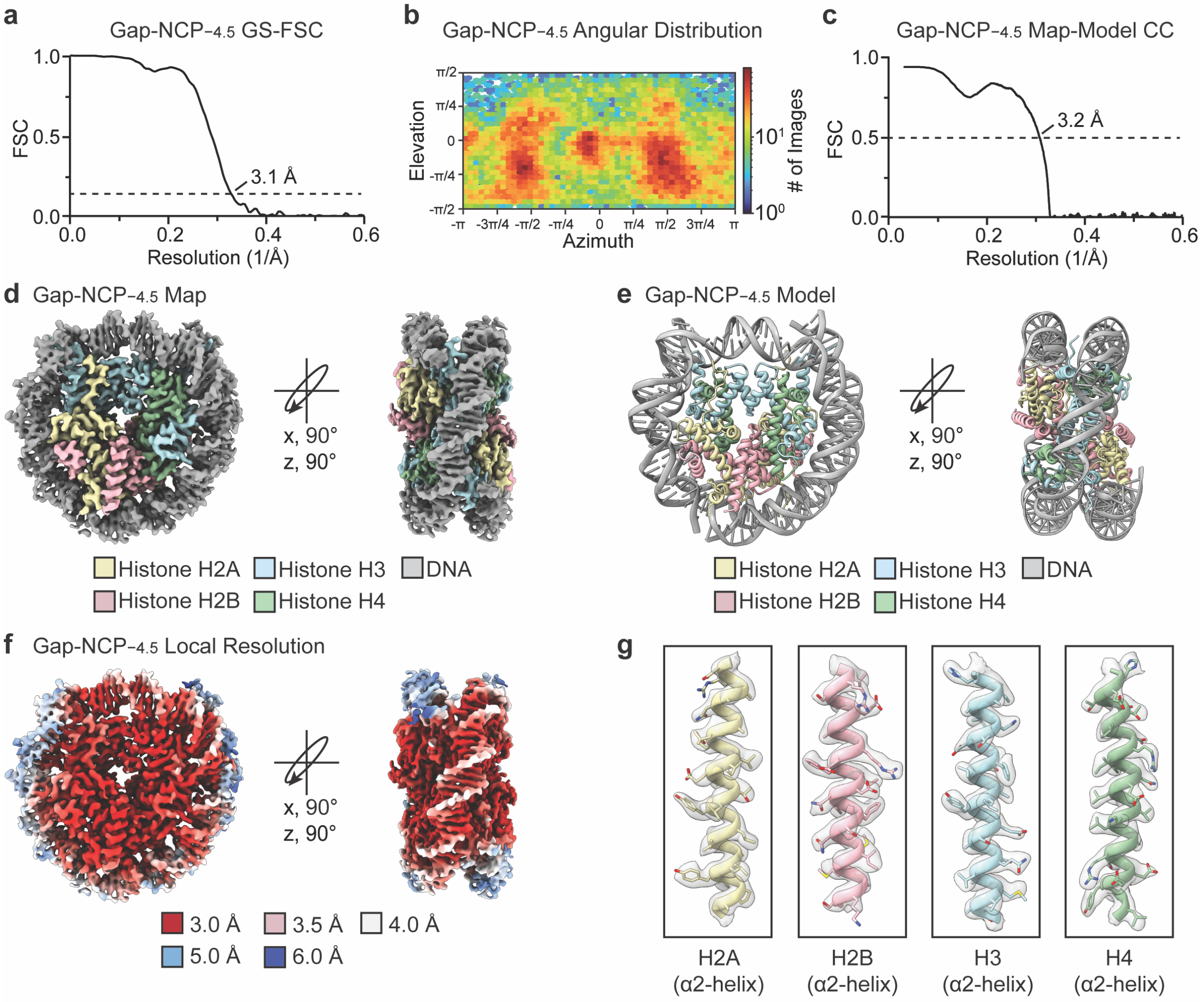

**Supplementary Fig. 4: Gap-NCP$_{-4.5}$ map and model quality assessment**

**a,** Gold-standard Fourier shell correlation (GS-FSC) curves for the Gap-NCP$_{-4.5}$ cryo-EM map (black line). The dashed line corresponds to GS-FSC - 0.143. **b,** Angular distribution heatmap for the Gap-NCP$_{-4.5}$ cryo-EM map**. c,** Map-to-model FSC curves for the Gap-NCP$_{-4.5}$ model and Gap-NCP$_{-4.5}$ cryo-EM map. The dashed line corresponds to FSC - 0.5. **d,** The final 3.1 Å Gap-NCP$_{-4.5}$ cryo-EM map shown in two different orientations. **e**, The final Gap-NCP$_{-4.5}$ model shown in two different orientations. **f,** The local resolution estimation for the Gap-NCP$_{-4.5}$ cryo-EM map shown in two different orientations. **g,** Representative segmented densities for histones H2A, H2B, H3, and H4 in the Gap-NCP$_{-4.5}$ cryo-EM map. The representative segmented densities from the cryo-EM map are shown as transparent gray surfaces.

#### Supplementary Fig. 5

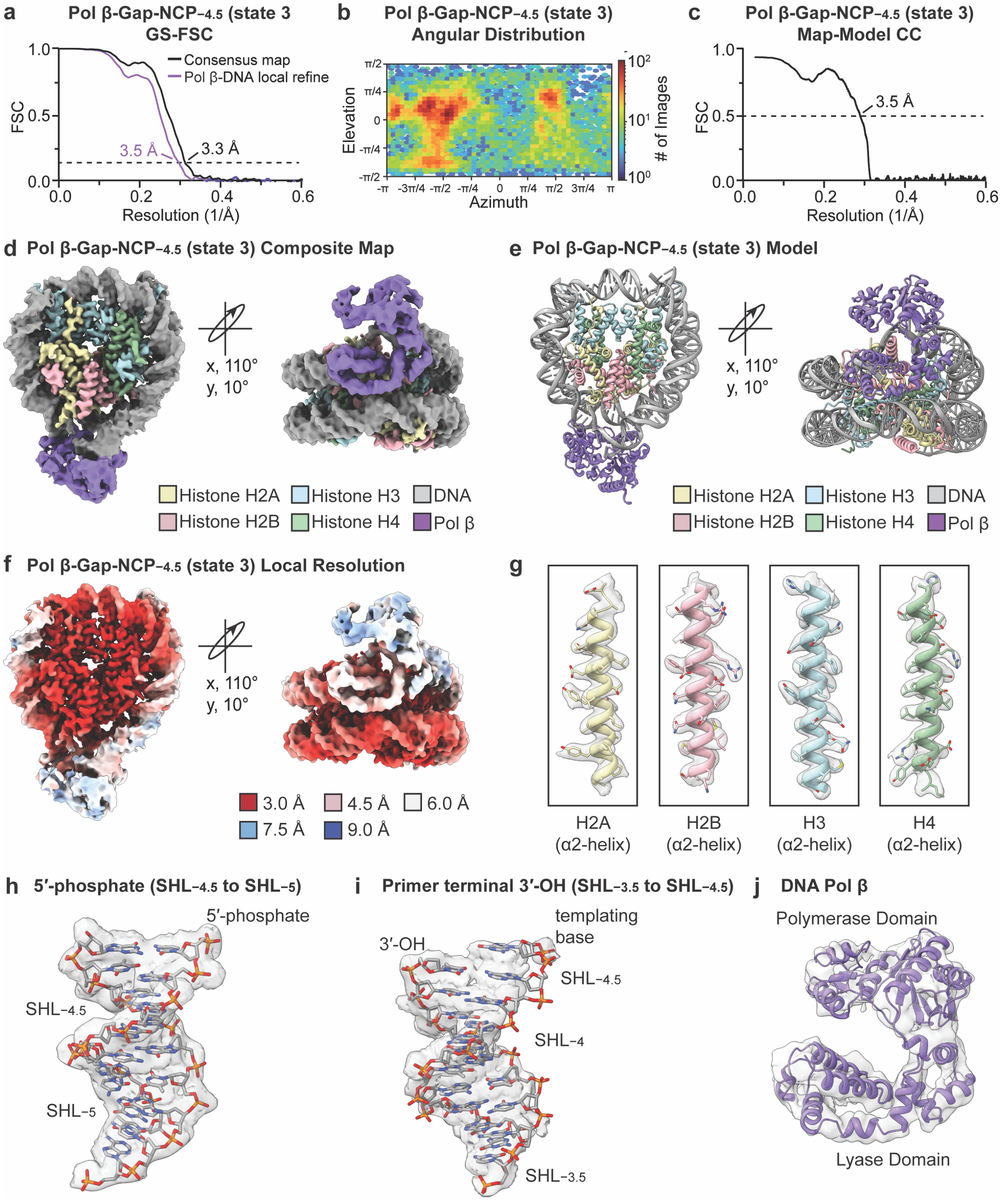

**Supplementary Fig. 5: Pre-catalytic Pol β-Gap-NCP$_{-4.5}$ map and model quality assessment**

**a,** Gold-standard Fourier shell correlation (GS-FSC) curves for the Pol β-Gap-NCP$_{-4.5}$ (pre-catalytic, state 3) consensus (black line) and Pol β/nucleosomal DNA focus (purple line) cryo-EM maps. The dashed line corresponds to GS-FSC - 0.143. **b,** Angular distribution heatmap for the Pol β-Gap-NCP$_{-4.5}$ (pre-catalytic, state 3) composite cryo-EM map**. c,** Map-to-model FSC curves for the Pol β-Gap-NCP$_{-4.5}$ (pre-catalytic, state 3) model and Pol β-Gap-NCP$_{-4.5}$ (pre-catalytic, state 3) composite cryo-EM map. The dashed line corresponds to FSC - 0.5. **d,** The final 3.3 Å Pol β-Gap-NCP$_{-4.5}$ (pre-catalytic, state 3) composite cryo-EM map shown in two different orientations. **e**, The Pol β-Gap-NCP$_{-4.5}$ (pre-catalytic, state 3) model shown in two different orientations. **f,** The local resolution estimation for the Pol β-Gap-NCP$_{-4.5}$ (pre-catalytic, state 3) composite cryo-EM map shown in two different orientations. **g,** Representative segmented densities for histones H2A, H2B, H3, and H4 in the Pol β-Gap-NCP$_{-4.5}$ (pre-catalytic, state 3) composite cryo-EM map. **h,** Representative segmented density for the 5′-phosphate and surrounding nucleosomal DNA from SHL$_{-4.5}$ to SHL$_{-5}$ in the Pol β-Gap-NCP$_{-4.5}$ (pre-catalytic, state 3) composite cryo-EM map. **I,** Representative segmented density for the primer terminal 3′-OH and surrounding nucleosomal DNA from SHL$_{-3.5}$ to SHL$_{-4.5}$ in the Pol β-Gap-NCP$_{-4.5}$ (pre-catalytic, state 3) composite cryo-EM map. **j,** Representative segmented density for Pol β in the Pol β-Gap-NCP$_{-4.5}$ (pre-catalytic, state 3) composite cryo-EM map. The representative segmented densities from the Pol β-Gap-NCP$_{-4.5}$ (pre-catalytic, state 3) in (g-j) are shown as transparent gray surfaces.

#### Supplementary Fig. 6

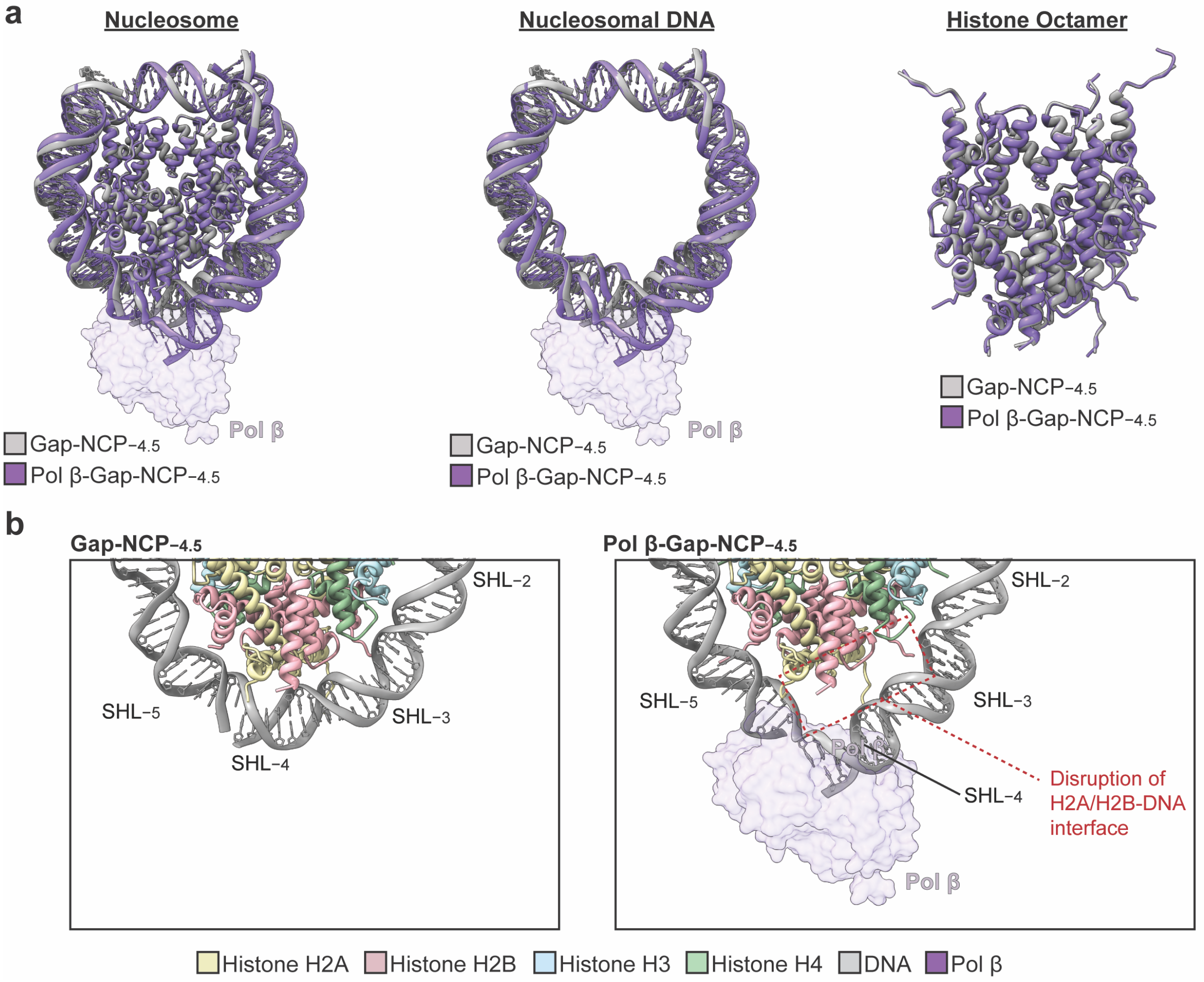

**Supplementary Fig. 6: Pol β induces structural distortions in the nucleosomal DNA during 1-nt gap recognition**

**a,** Structural comparison of the nucleosome (left), nucleosomal DNA (middle), and histone octamer (right) in the Gap-NCP$_{-4.5}$ (gray) and the pre-catalytic Pol β-Gap-NCP$_{-4.5}$ complex (purple). **b,** Focused views of the nucleosomal DNA from SHL−2.5 to SHL−5.5 in the Gap-NCP$_{-4.5}$ (left) and Pol β-Gap-NCP$_{-4.5}$ complex (right). Pol β is shown as a transparent purple surface in (a-b).

#### Supplementary Fig. 7

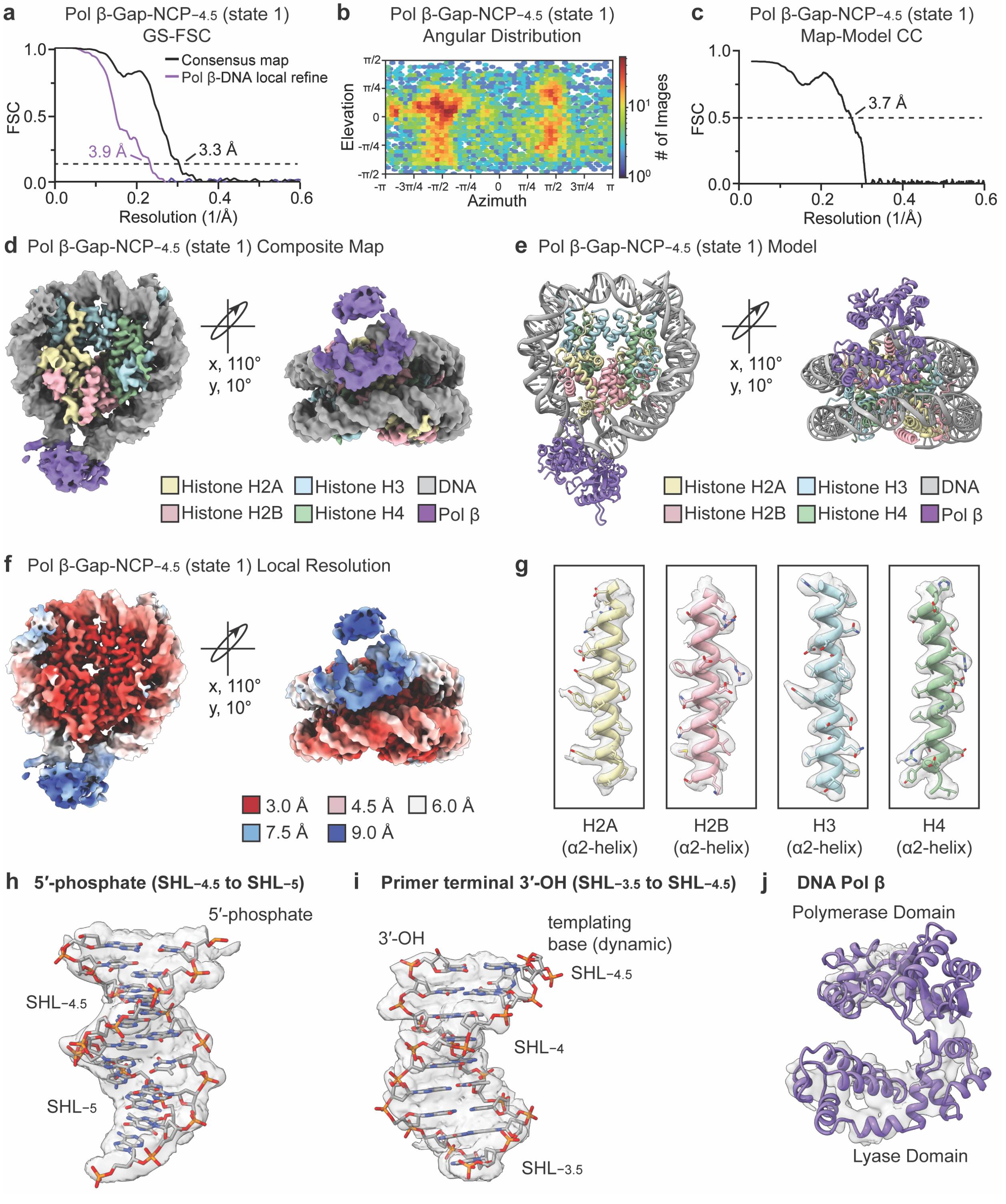

**Supplementary Fig. 7: Pol β-Gap-NCP−4.5 (5′-phosphate capture, state 1) map and model quality assessment**

**a,** Gold-standard Fourier shell correlation (GS-FSC) curves for the Pol β-Gap-NCP−4.5 (5′-phosphate capture, state 1) consensus (black line) and Pol β/nucleosomal DNA focus (purple line) cryo-EM maps. The dashed line corresponds to GS-FSC - 0.143. **b,** Angular distribution heatmap for the Pol β-Gap-NCP−4.5 (5′-phosphate capture, state 1) composite cryo-EM map**. c,** Map-to-model FSC curves for the Pol β-Gap-NCP−4.5 (5′-phosphate capture, state 1) model and Pol β-Gap-NCP−4.5 (5′-phosphate capture, state 1) composite cryo-EM map. The dashed line corresponds to FSC - 0.5. **d,** The final 3.3 Å Pol β-Gap-NCP−4.5 (5′-phosphate capture, state 1) composite cryo-EM map shown in two different orientations. **e**, The Pol β-Gap-NCP−4.5 (5′-phosphate capture, state 1) model shown in two different orientations. **f,** The local resolution estimation for the Pol β-Gap-NCP−4.5 (5′-phosphate capture, state 1) composite cryo-EM map shown in two different orientations. **g,** Representative segmented densities for histones H2A, H2B, H3, and H4 in the Pol β-Gap-NCP−4.5 (5′-phosphate capture, state 1) composite cryo-EM map. **h,** Representative segmented density for the 5′-phosphate and surrounding nucleosomal DNA from SHL−4.5 to SHL−5 in the Pol β-Gap-NCP−4.5 (5′-phosphate capture, state 1) composite cryo-EM map. **I,** Representative segmented density for the primer terminal 3′-OH and surrounding nucleosomal DNA from SHL−3.5 to SHL−4.5 in the Pol β-Gap-NCP−4.5 (5′-phosphate capture, state 1) composite cryo-EM map. **j,** Representative segmented density for Pol β in the Pol β-Gap-NCP−4.5 (5′-phosphate capture, state 1) composite cryo-EM map. The representative segmented densities from the Pol β-Gap-NCP−4.5 (5′-phosphate capture, state 1) in (g-j) are shown as transparent gray surfaces.

#### Supplementary Fig. 8

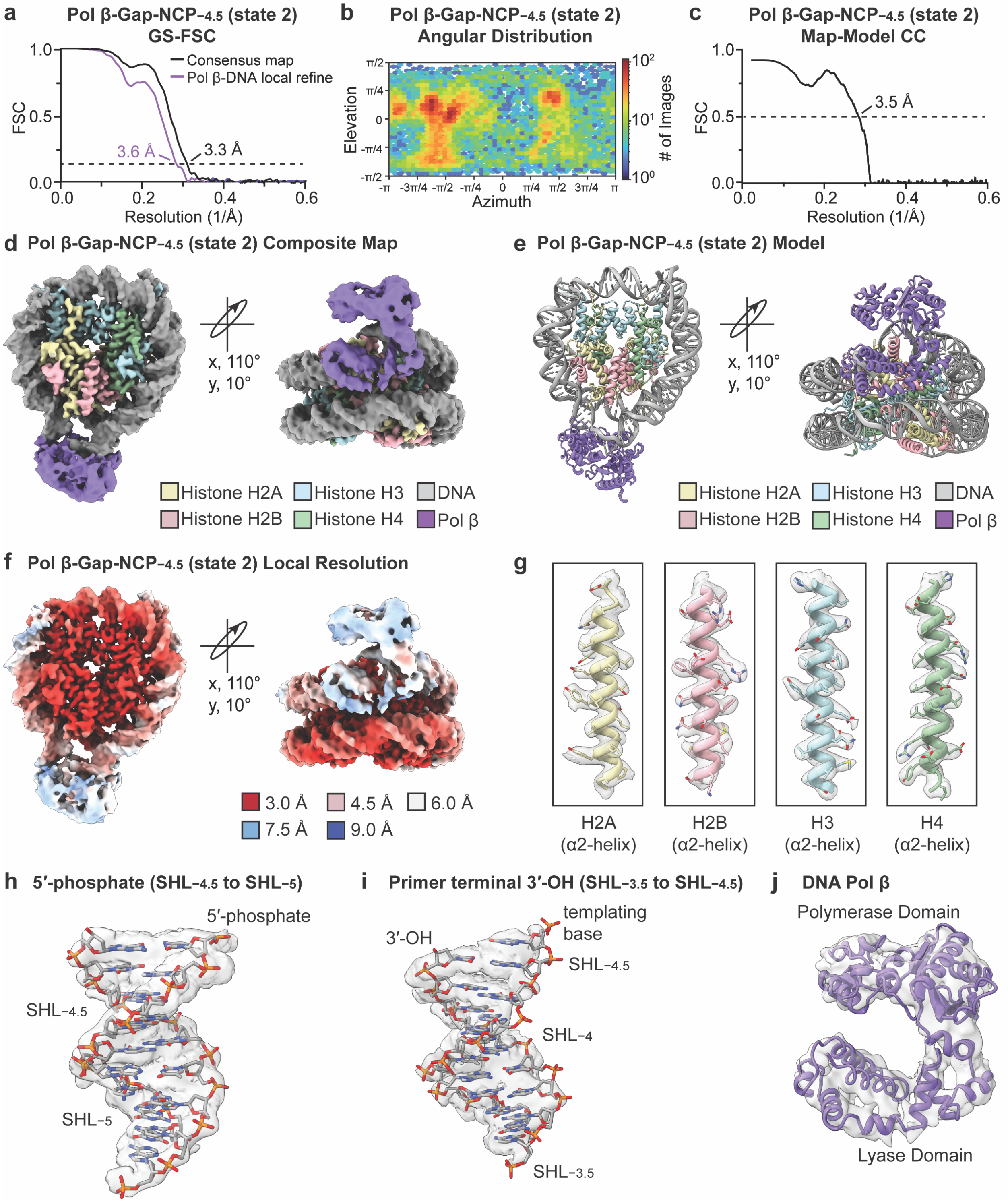

**Supplementary Fig. 8: Pol β-Gap-NCP$_{-4.5}$ (primer terminus capture, state 2) map and model quality assessment**

**a,** Gold-standard Fourier shell correlation (GS-FSC) curves for the Pol β-Gap-NCP$_{-4.5}$ (primer terminus capture, state 2) consensus (black line) and Pol β/nucleosomal DNA focus (purple line) cryo-EM maps. The dashed line corresponds to GS-FSC - 0.143. **b,** Angular distribution heatmap for the Pol β-Gap-NCP$_{-4.5}$ (primer terminus capture, state 2) composite cryo-EM map**. c,** Map-to-model FSC curves for the Pol β-Gap-NCP$_{-4.5}$ (primer terminus capture, state 2) model and Pol β-Gap-NCP$_{-4.5}$ (primer terminus capture, state 2) composite cryo-EM map. The dashed line corresponds to FSC - 0.5. **d,** The final 3.3 Å Pol β-Gap-NCP$_{-4.5}$ (primer terminus capture, state 2) composite cryo-EM map shown in two different orientations. **e**, The Pol β-Gap-NCP$_{-4.5}$ (primer terminus capture, state 2) model shown in two different orientations. **f,** The local resolution estimation for the Pol β-Gap-NCP$_{-4.5}$ (primer terminus capture, state 2) composite cryo-EM map shown in two different orientations. **g,** Representative segmented densities for histones H2A, H2B, H3, and H4 in the Pol β-Gap-NCP$_{-4.5}$ (primer terminus capture, state 2) composite cryo-EM map. **h,** Representative segmented density for the 5′-phosphate and surrounding nucleosomal DNA from SHL$_{-4.5}$ to SHL$_{-5}$ in the Pol β-Gap-NCP$_{-4.5}$ (primer terminus capture, state 2) composite cryo-EM map. **I,** Representative segmented density for the primer terminal 3′-OH and surrounding nucleosomal DNA from SHL$_{-3.5}$ to SHL$_{-4.5}$ in the Pol β-Gap-NCP$_{-4.5}$ (primer terminus capture, state 2) composite cryo-EM map. **j,** Representative segmented density for Pol β in the Pol β-Gap-NCP$_{-4.5}$ (primer terminus capture, state 2) composite cryo-EM map. The representative segmented densities from the Pol β-Gap-NCP$_{-4.5}$ (primer terminus capture, state 2) in (g-j) are shown as transparent gray surfaces.

**Supplementary Fig. 9**

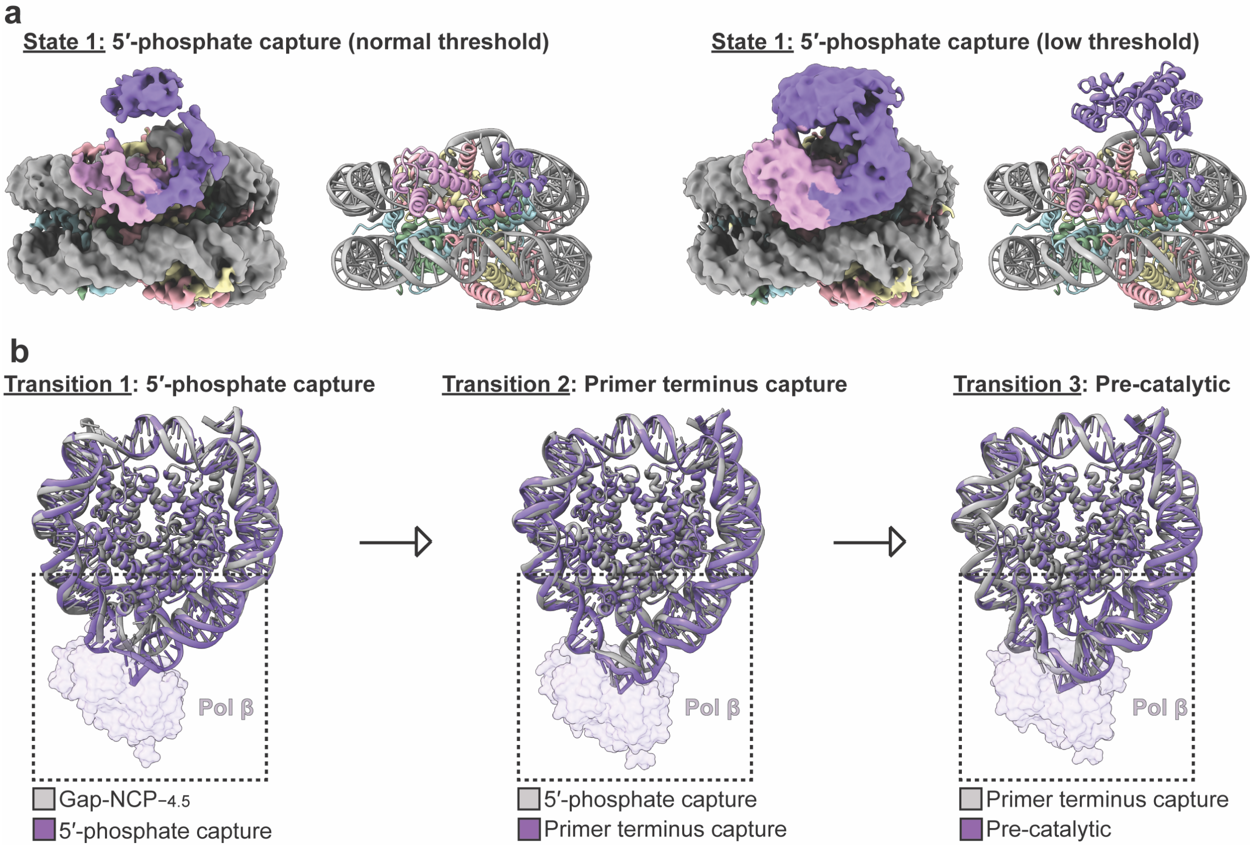

**Supplementary Fig. 9: Structural analysis of the Pol β-Gap-NCP$_{-4.5}$ states**

**a,** The Pol β-Gap-NCP$_{-4.5}$ (5′-phosphate capture, state 1) composite cryo-EM map and model (Pol β residues 11-146). The Pol β-Gap-NCP$_{-4.5}$ (5′-phosphate capture, state 1) composite cryo-EM map is shown at normal threshold. **b,** The Pol β-Gap-NCP$_{-4.5}$ (5′-phosphate capture, state 1) composite cryo-EM map and model. The Pol β-Gap-NCP$_{-4.5}$ (5′-phosphate capture, state 1) composite cryo-EM map is shown at low threshold. **c,** Structural comparison of the nucleosomal DNA during each individual transition from the apo Gap-NCP$_{-4.5}$ to the pre-catalytic state (state 3). Pol β is shown as a transparent purple surface.

**Supplementary Fig. 10**

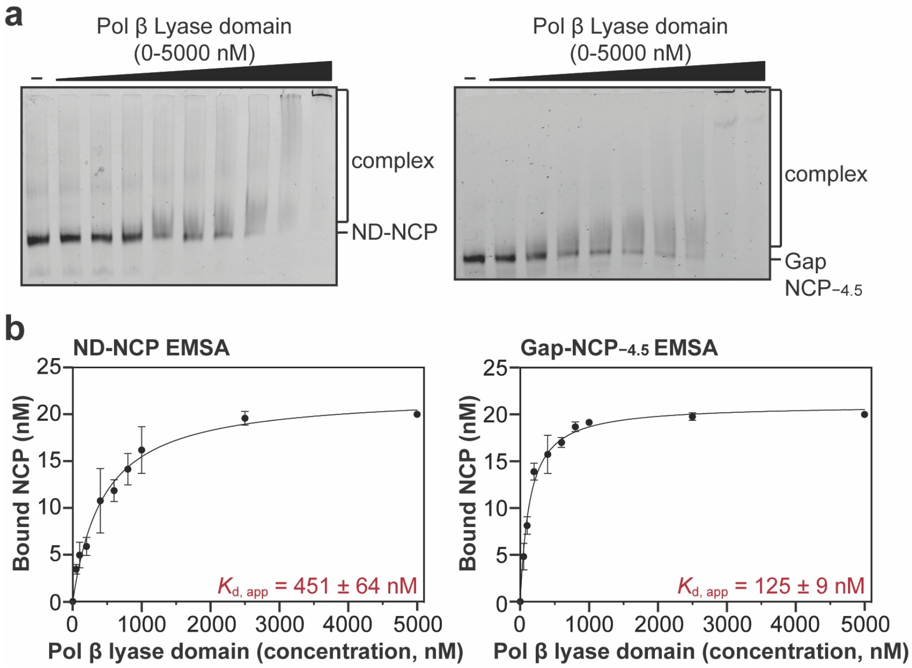

**Supplementary Fig. 10: Analysis of nucleosome binding by the Pol β lyase domain**

**a,** Representative native PAGE gels from electrophoretic mobility shift assays (EMSAs) of Pol β lyase domain (residues 1-87) and a ND-NCP and Gap-NCP$_{-4.5}$. These gels are representative of three independent replicate EMSA experiments for each NCP. **b,** Quantification and fits from the EMSA experiments with the Pol β lyase domain (residues 1-87) and a ND-NCP and Gap-NCP$_{-4.5}$. The data points represent the mean ± standard deviation from three independent replicate experiments. The error bars are included for all experimental data points, but some error bars are smaller than the circles used to represent the data points. The apparent binding affinity ($K_{d, app}$) is shown as an inset for each experiment and represents the mean ± standard error of the mean from the three independent replicate experiments. All source data in this figure are provided as a Source Data file.

**Supplementary Fig. 11**

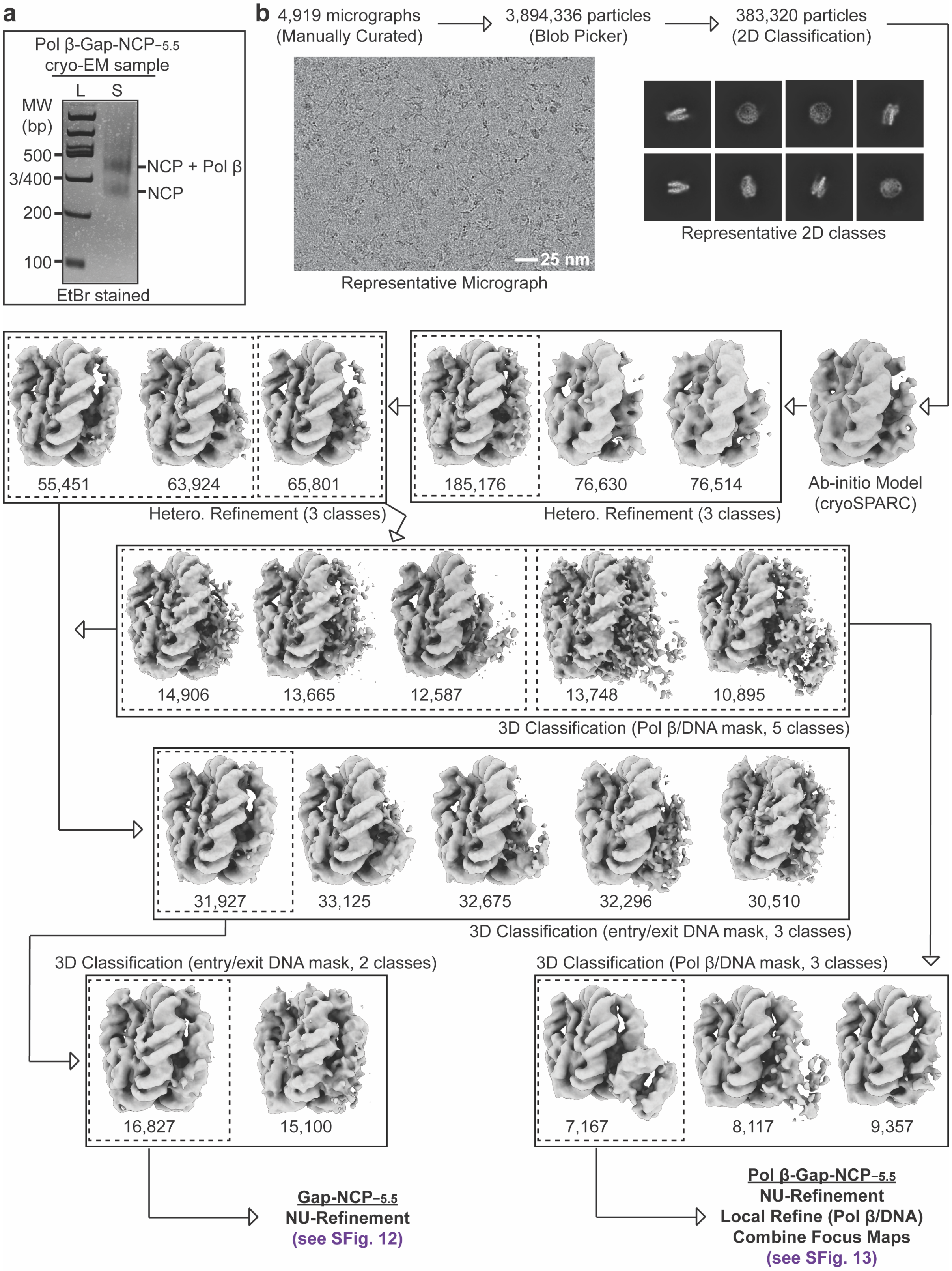

**Supplementary Fig. 11: Pol β-Gap-NCP$_{-5.5}$ SPA processing workflow**

**a,** Native PAGE gel of the Pol β-Gap-NCP$_{-5.5}$ cryo-EM sample (S) and a 100 bp DNA ladder (L). The Gap-NCP$_{-5.5}$ and Pol β-Gap-NCP$_{-5.5}$ complex were visualized with ethidium bromide staining. **b,** Flowchart of the data processing pipeline for the Pol β-Gap-NCP$_{-5.5}$ cryo-EM dataset. A representative micrograph (n=4,919) and representative 2D classes from the Pol β-Gap-NCP$_{-5.5}$ cryo-EM dataset are shown. The maps chosen for further classification and/or refinement throughout the data processing pipeline are boxed. The final maps, final models, and quality assessment metrics for Gap-NCP$_{-5.5}$ and the pre-catalytic Pol β-Gap-NCP$_{-5.5}$ complex can be found in Supplementary Figs. 12 and 13.

#### Supplementary Fig. 12

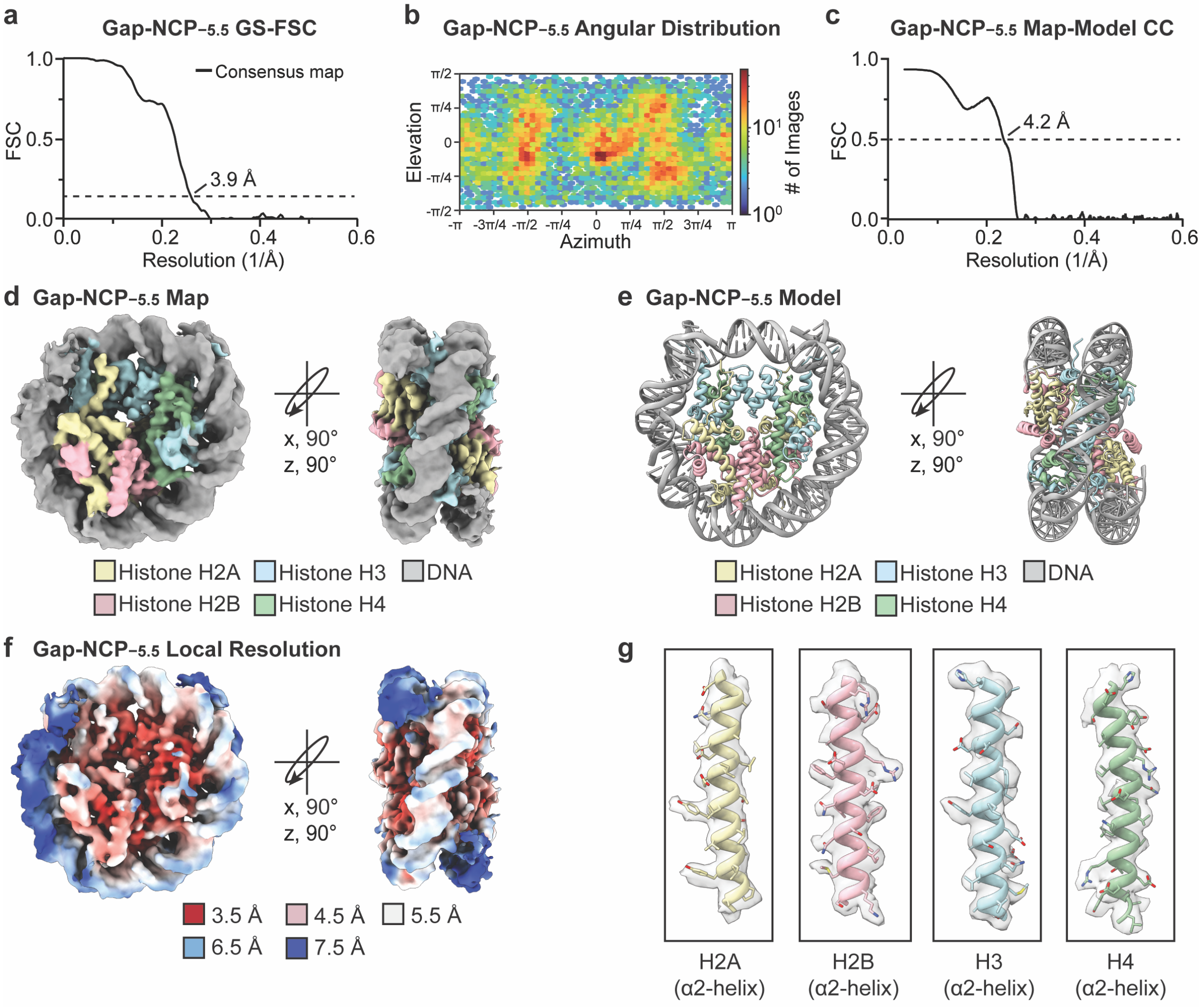

**Supplementary Fig. 12: Gap-NCP$_{-5.5}$ map and model quality assessment**

**a,** Gold-standard Fourier shell correlation (GS-FSC) curves for the Gap-NCP$_{-5.5}$ cryo-EM map (black line). The dashed line corresponds to GS-FSC - 0.143. **b,** Angular distribution heatmap for the Gap-NCP$_{-5.5}$ cryo-EM map**. c,** Map-to-model FSC curves for the Gap-NCP$_{-5.5}$ model and Gap-NCP$_{-5.5}$ cryo-EM map. The dashed line corresponds to FSC - 0.5. **d,** The final 3.9 Å Gap-NCP$_{-3.5}$ cryo-EM map shown in two different orientations. **e**, The final Gap-NCP$_{-5.5}$ model shown in two different orientations. **f,** The local resolution estimation for the Gap-NCP$_{-5.5}$ cryo-EM map shown in two different orientations. **g,** Representative segmented densities for histones H2A, H2B, H3, and H4 in the Gap-NCP$_{-5.5}$ cryo-EM map. The representative segmented densities from the cryo-EM map are shown as transparent gray surfaces.

#### Supplementary Fig. 13

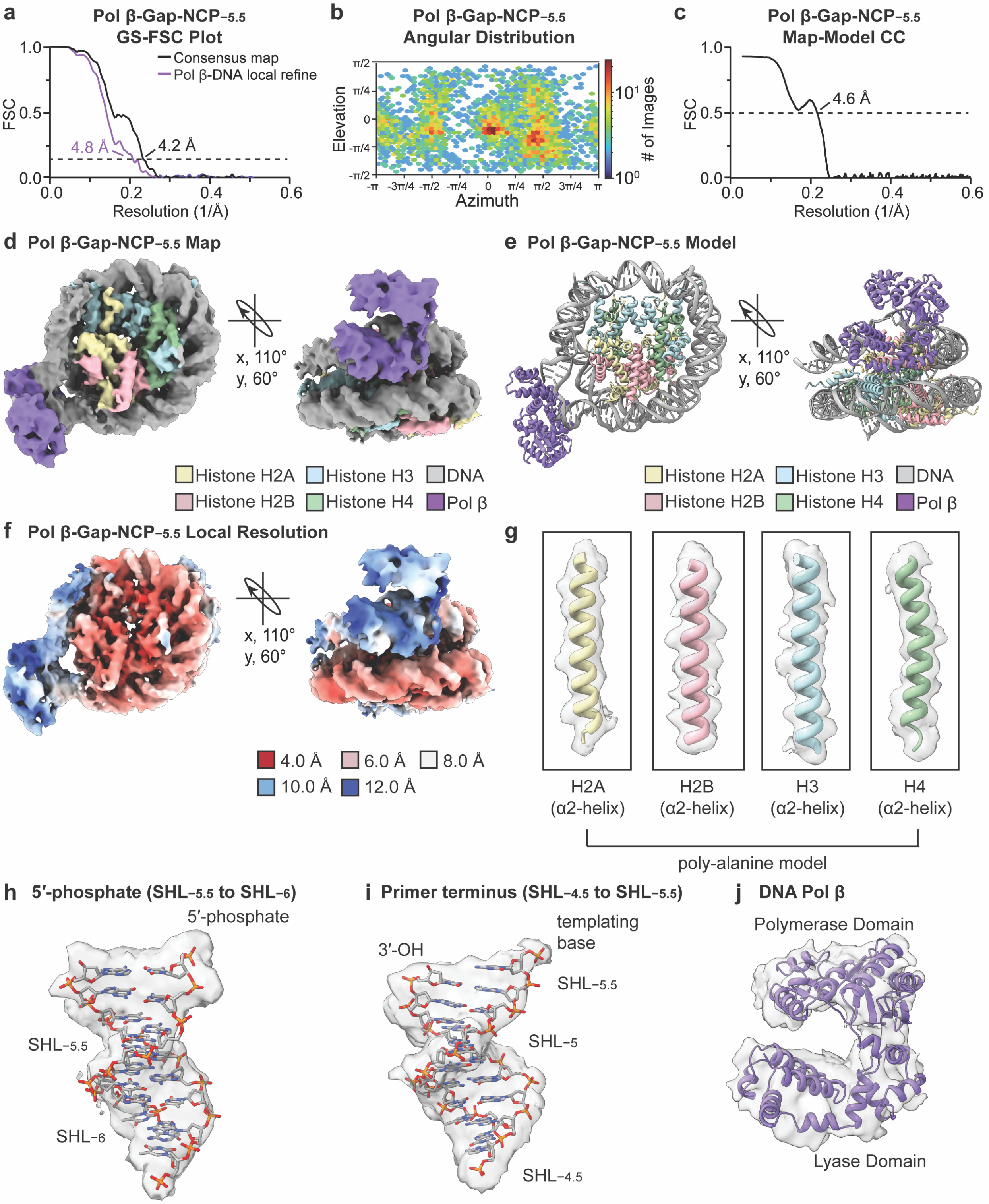

**Supplementary Fig. 13: Pol β-Gap-NCP$_{-5.5}$ map and model quality assessment**

**a,** Gold-standard Fourier shell correlation (GS-FSC) curves for the Pol β-Gap-NCP$_{-5.5}$ consensus (black line) and Pol β/nucleosomal DNA focus (purple line) cryo-EM maps. The dashed line corresponds to GS-FSC - 0.143. **b,** Angular distribution heatmap for the Pol β-Gap-NCP$_{-5.5}$ composite cryo-EM map**. c,** Map-to-model FSC curves for the Pol β-Gap-NCP$_{-5.5}$ model and Pol β-Gap-NCP$_{-5.5}$ composite cryo-EM map. The dashed line corresponds to FSC - 0.5. **d,** The final 4.2 Å Pol β-Gap-NCP$_{-5.5}$ composite cryo-EM map shown in two different orientations. **e**, The Pol β-Gap-NCP$_{-5.5}$ model shown in two different orientations. **f,** The local resolution estimation for the Pol β-Gap-NCP$_{-5.5}$ composite cryo-EM map shown in two different orientations. **g,** Representative segmented densities for histones H2A, H2B, H3, and H4 in the Pol β-Gap-NCP$_{-5.5}$ composite cryo-EM map. **h,** Representative segmented density for the 5′-phosphate and surrounding nucleosomal DNA from SHL$_{-5.5}$ to SHL$_{-6}$ in the Pol β-Gap-NCP$_{-5.5}$ composite cryo-EM map. **I,** Representative segmented density for the primer terminal 3′-OH and surrounding nucleosomal DNA from SHL$_{-4.5}$ to SHL$_{-5.5}$ in the Pol β-Gap-NCP$_{-5.5}$ composite cryo-EM map. **j,** Representative segmented density for Pol β in the Pol β-Gap-NCP$_{-5.5}$ composite cryo-EM map. The representative segmented densities from the Pol β-Gap-NCP$_{-5.5}$ in (g-j) are shown as transparent gray surfaces.

#### Supplementary Fig. 14

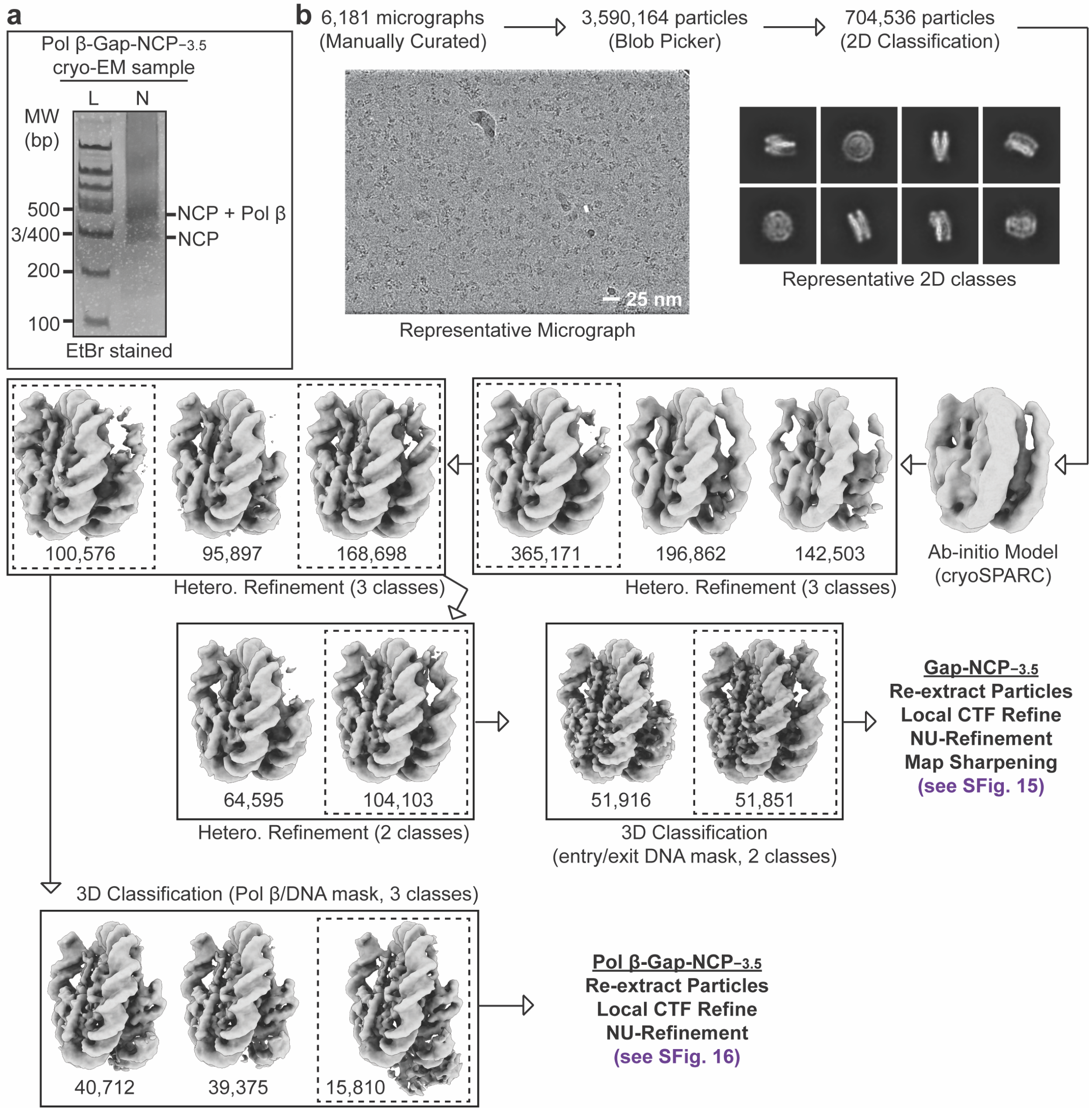

**Supplementary Fig. 14: Pol β-Gap-NCP$_{-3.5}$ SPA processing workflow**

**a,** Native PAGE gel of the Pol β-Gap-NCP$_{-3.5}$ cryo-EM sample (S) and a 100 bp DNA ladder (L). The Gap-NCP$_{-3.5}$ and Pol β-Gap-NCP$_{-3.5}$ complex were visualized with ethidium bromide staining. **b,** Flowchart of the data processing pipeline for the Pol β-Gap-NCP$_{-3.5}$ cryo-EM dataset. A representative micrograph (n=6181) and representative 2D classes from the Pol β-Gap-NCP$_{-3.5}$ cryo-EM dataset are shown. The maps chosen for further classification and/or refinement throughout the data processing pipeline are boxed. The final maps, final models, and quality assessment metrics for Gap-NCP$_{-3.5}$ and Pol β-Gap-NCP$_{-3.5}$ complex can be found in Supplementary Figs. 15 and 16.

#### Supplementary Fig. 15

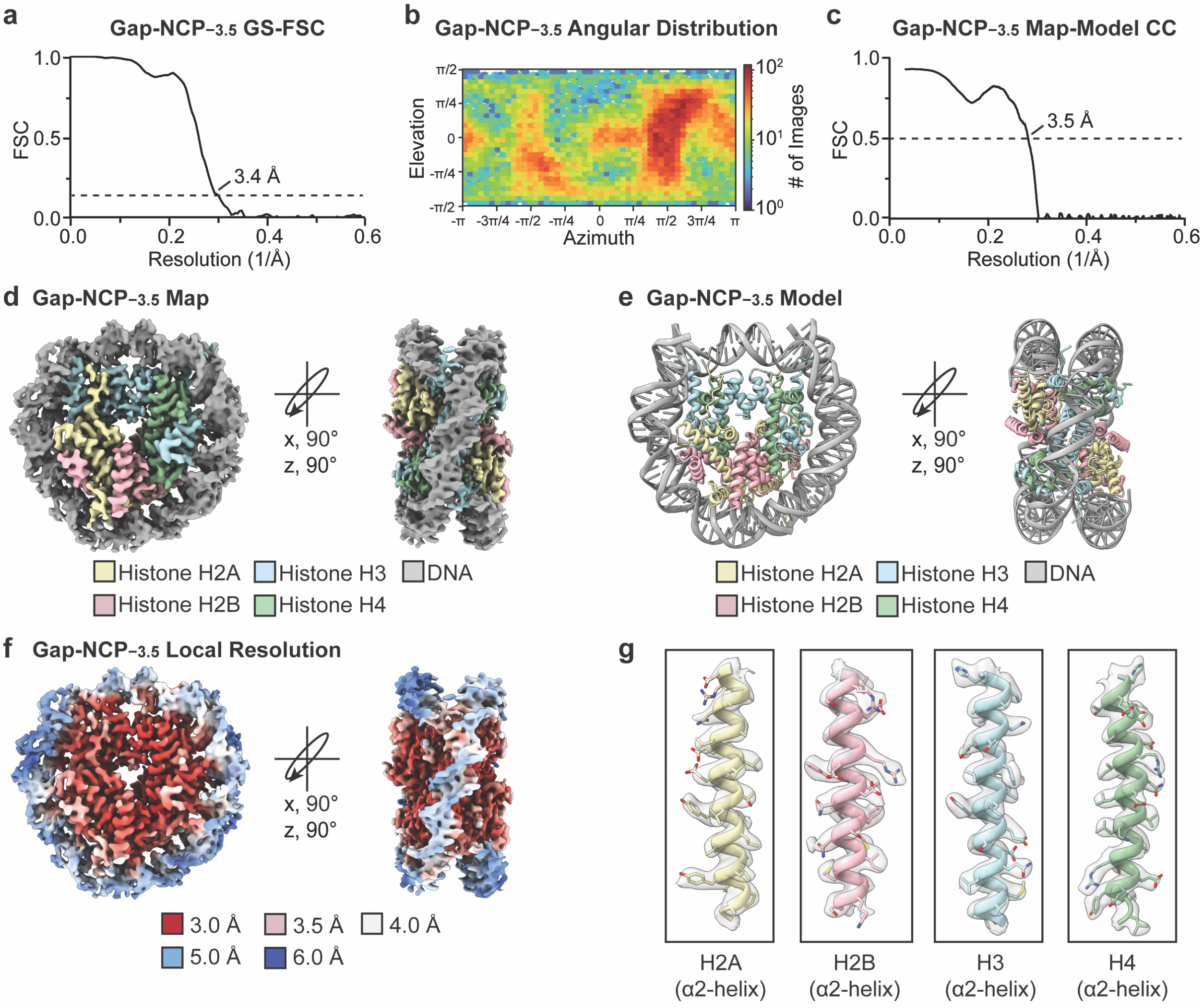

**Supplementary Fig. 15: Gap-NCP$_{-3.5}$ map and model quality assessment**

**a,** Gold-standard Fourier shell correlation (GS-FSC) curves for the Gap-NCP$_{-3.5}$ cryo-EM map (black line). The dashed line corresponds to GS-FSC - 0.143. **b,** Angular distribution heatmap for the Gap-NCP$_{-3.5}$ cryo-EM map**. c,** Map-to-model FSC curves for the Gap-NCP$_{-3.5}$ model and Gap-NCP$_{-3.5}$ cryo-EM map. The dashed line corresponds to FSC - 0.5. **d,** The final 3.4 Å Gap-NCP$_{-3.5}$ cryo-EM map shown in two different orientations. **e**, The final Gap-NCP$_{-3.5}$ model shown in two different orientations. **f,** The local resolution estimation for the Gap-NCP$_{-3.5}$ cryo-EM map shown in two different orientations. **g,** Representative segmented densities for histones H2A, H2B, H3, and H4 in the Gap-NCP$_{-3.5}$ cryo-EM map. The representative segmented densities from the cryo-EM map are shown as transparent gray surfaces.

#### Supplementary Fig. 16

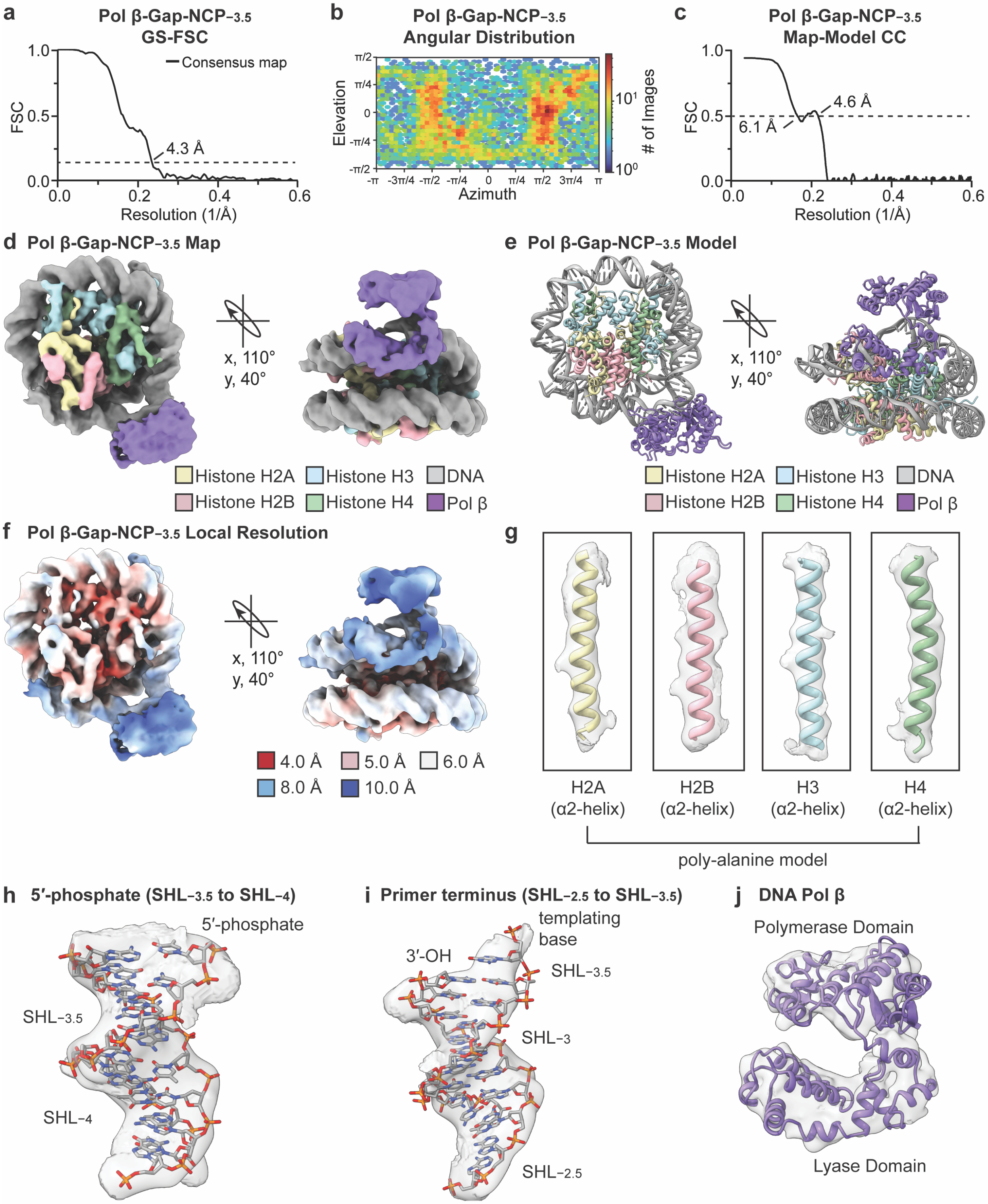

**Supplementary Fig. 16: Pol β-Gap-NCP$_{-3.5}$ map and model quality assessment**

**a,** Gold-standard Fourier shell correlation (GS-FSC) curves for the Pol β-Gap-NCP$_{-3.5}$ cryo-EM map (black line). The dashed line corresponds to GS-FSC - 0.143. **b,** Angular distribution heatmap for the Pol β-Gap-NCP$_{-3.5}$ cryo-EM map**. c,** Map-to-model FSC curves for the Pol β-Gap-NCP$_{-3.5}$ cryo-EM map. The dashed line corresponds to FSC - 0.5. **d,** The final 4.3 Å Pol β-Gap-NCP$_{-3.5}$ cryo-EM map shown in two different orientations. **e**, The Pol β-Gap-NCP$_{-3.5}$ model shown in two different orientations. **f,** The local resolution estimation for the Pol β-Gap-NCP$_{-3.5}$ cryo-EM map shown in two different orientations. **g,** Representative segmented densities for histones H2A, H2B, H3, and H4 in the Pol β-Gap-NCP$_{-3.5}$ cryo-EM map. **h,** Representative segmented density for the 5′-phosphate and surrounding nucleosomal DNA from SHL$_{-3.5}$ to SHL$_{-4}$ in the Pol β-Gap-NCP$_{-3.5}$ cryo-EM map. **I,** Representative segmented density for the primer terminal 3′-OH and surrounding nucleosomal DNA from SHL$_{-2.5}$ to SHL$_{-3.5}$ in the Pol β-Gap-NCP$_{-3.5}$ cryo-EM map. **j,** Representative segmented density for Pol β in the Pol β-Gap-NCP$_{-3.5}$ cryo-EM map. The representative segmented densities from the Pol β-Gap-NCP$_{-3.5}$ cryo-EM map in (g-j) are shown as transparent gray surfaces.

**Supplementary Table 1**

| Substrate | Template | dNTP | $k_{obs}$ ($s^{-1}$) | Fold Change | Product (%) | $K_{d,app}$ (nM) | Fold Change |
|---|---|---|---|---|---|---|---|
| Gap DNA | dG | dCTP | 0.30 ± 0.02 | - | 82 ± 2 | 30 ± 10* | - |
| ND-NCP | - | - | - | - | - | 217 ± 14 | - |
| Gap-NCP$_{-5.5}$ | dC | dGTP | 0.12 ± $2.7 \times 10^{-3}$ | 2.5x | 83 ± 3 | 51 ± 3 | 4.3 |
| Gap-NCP$_{-4.5}$ | dG | dCTP | 0.01 ± $1 \times 10^{-3}$ | 33x | 89 ± 1 | 98 ± 4 | 2.2 |
| Gap-NCP$_{-3.5}$ | dG | dCTP | $4.6 \times 10^{-3} \pm 1.6 \times 10^{-4}$ | 65x | 73 ± 1 | 104 ± 19 | 2.1 |
| Gap-NCP$_{-2.5}$ | dG | dCTP | $1.8 \times 10^{-3} \pm 1.4 \times 10^{-4}$ | 167x | 88 ± 3 | 101 ± 16 | 2.1 |
| Gap-NCP$_{-1.5}$ | dA | dTTP | - | - | 10 | 133 ± 20 | 1.6 |
| Gap-NCP$_{-4.5+2}$ | dG | dCTP | $4.7 \times 10^{-4} \pm 2.0 \times 10^{-4}$ | 638x | 61 ± 4 | 143 ± 9 | 1.5 |
| Gap-NCP$_{-4.5+3}$ | dA | dTTP | $3.9 \times 10^{-4} \pm 1.9 \times 10^{-4}$ | 770x | 26 ± 2 | 137 ± 11 | 1.6 |
| Gap-NCP$_{-4.5+4}$ | dC | dGTP | $4.9 \times 10^{-4} \pm 5.1 \times 10^{-5}$ | 612x | 31 ± 3 | 104 ± 5 | 2.1 |

*$K_{d,app}$ value from Howard et al. *JBC*, 2020

**Supplementary Table 2**

| Data collection and processing | | | | |
|---|---|---|---|---|
| **Dataset** | **Gap-NCP−4.5** | | | |
| Magnification | 105,000 | | | |
| Voltage (kV) | 300 | | | |
| Electron exposure ($e^{-}/Å^{2}$) | 50 | | | |
| Defocus range (μm) | −0.8 to −2.2 | | | |
| Pixel size (Å) | 0.413 | | | |
| Symmetry imposed | C1 | | | |
| Initial particle images (no.) | 3,908,068 | | | |
| **Structure** | **Gap-NCP−4.5** | **Polβ-Gap-NCP−4.5 (5′ capture, state 1)** | **Polβ-Gap-NCP−4.5 (PT capture, state 2)** | **Polβ-Gap-NCP−4.5 (pre-catalytic, state 3)** |
| Final particle images (no.) | 42,803 | 17,090 | 27,265 | 29,841 |
| Map resolution (Å) | 3.1 | 3.3 | 3.3 | 3.3 |
| FSC threshold | 0.143 | 0.143 | 0.143 | 0.143 |
| PDB accession | **9DWF** | **9DWG** | **9DWH** | **9DWI** |
| EMDB accession | **EMD-47242** | **EMD-47243<br>EMD-47244<br>EMD-47245** | **EMD-47246<br>EMD-47247<br>EMD-47248** | **EMD-47249<br>EMD-47250<br>EMD-47251** |
| **Refinement** | | | | |
| Initial model used (PDB ID) | 7U52 | 7U52, 3ISB | 7U52, 3ISB | 7U52, 3ISB |
| Model resolution (Å) | 3.2 | 3.7 | 3.5 | 3.5 |
| FSC threshold | 0.5 | 0.5 | 0.5 | 0.5 |
| **Model composition** | | | | |
| Nonhydrogen atoms | 11,948 | 14,535 | 14,653 | 14,661 |
| Protein residues | 749 | 1074 | 1085 | 1085 |
| Nucleotide | 293 | 293 | 293 | 293 |
| **B factors ($Å^2$ )** | | | | |
| Protein | 24.12 | 254.39 | 132.74 | 134.54 |
| Nucleotide | 76.67 | 238.55 | 158.33 | 158.87 |
| **r.m.s. deviations** | | | | |
| Bond Length (Å) (# > 4σ) | 0.004 0) | 0.006 (4) | 0.004 (1) | 0.004 (1) |
| Bond Angles (º) (# > 4σ) | 0.583 (1) | 0.947 (7) | 0.656 (15) | 0.634 (4) |
| **Validation** | | | | |
| MolProbity score | 1.21 | 1.66 | 1.32 | 1.46 |
| Clashscore | 4.28 | 7.91 | 5.57 | 5.78 |
| Poor rotamers (%) | 2.09 | 0.89 | 0.22 | 0.11 |
| **Ramachandran plot** | | | | |
| Favored (%) | 98.77 | 96.49 | 97.93 | 97.18 |
| Allowed (%) | 1.23 | 3.51 | 2.07 | 2.82 |
| Disallowed (%) | 0.00 | 0.00 | 0.00 | 0.00 |

**Supplementary Table 3**

| **Data collection and processing** | | | | |
|---|---|---|---|---|
| **Dataset** | **Gap-NCP$^{-5.5}$** | | **Gap-NCP$^{-3.5}$** | |
| Magnification | 130,000 | | 29,000 | |
| Voltage (kV) | 300 | | 300 | |
| Electron exposure ($e^{-}/Å^{2}$) | 60 | | 51 | |
| Defocus range (µm) | -0.5 to -2.5 | | −0.8 to −2.4 | |
| Pixel size (Å) | 0.970 | | 0.394 | |
| Symmetry imposed | C1 | | C1 | |
| Initial particle images (no.) | 3,382,261 | | 3,590,164 | |
| **Structure** | **Gap-NCP$^{-5.5}$** | **Polβ-Gap-NCP$^{-5.5}$ (poly-alanine)** | **Gap-NCP$^{-3.5}$** | **Polβ-Gap-NCP$^{-3.5}$ (poly-alanine)** |
| Final particle images (no.) | 16,827 | 7,167 | 51,851 | 15,757 |
| Map resolution (Å) | 3.9 | 4.2 | 3.4 | 4.3 |
| FSC threshold | 0.143 | 0.143 | 0.143 | 0.143 |
| PDB accession | **9DWL** | **9DWM** | **9DWJ** | **9DWK** |
| EMDB accession | **EMD-47254** | **EMD-47255**<br>**EMD-47256**<br>**EMD-47257** | **EMD-47252** | **EMD-47253** |
| **Refinement** | | | | |
| Initial model used (PDB ID) | 7U52 | 7U52, 3ISB | 7U52 | 7U52, 3ISB |
| Model resolution (Å) | 4.2 | 4.6 | 3.5 | 4.6 |
| FSC threshold | 0.5 | 0.5 | 0.5 | 0.5 |
| **Model composition** | | | | |
| Nonhydrogen atoms | 11,877 | 11,175 | 11,887 | 10,375 |
| Protein residues | 742 | 1065 | 748 | 1068 |
| Nucleotide | 293 | 289 | 291 | 249 |
| **B factors ($Å^2$)** | | | | |
| Protein | 150.12 | 405.63 | 48.09 | 381.79 |
| Nucleotide | 256.25 | 521.68 | 114.64 | 379.95 |
| **r.m.s. deviations** | | | | |
| Bond Length (Å) (# > 4σ) | 0.005 (1) | 0.005 (1) | 0.005 (1) | 0.005 (1) |
| Bond Angles (°) (# > 4σ) | 0.725 (4) | 0.612 (0) | 0.699 (8) | 0.629 (1) |
| **Validation** | | | | |
| MolProbity score | 1.39 | 1.54 | 1.49 | 1.41 |
| Clashscore | 7.03 | 6.64 | 5.75 | 7.56 |
| Poor rotamers (%) | 0.65 | 0.00 | 0.64 | 0.00 |
| **Ramachandran plot** | | | | |
| Favored (%) | 98.35 | 97.03 | 96.99 | 98.95 |
| Allowed (%) | 1.65 | 2.97 | 3.01 | 1.05 |
| Disallowed (%) | 0.00 | 0.00 | 0.00 | 0.00 |

**Supplementary Table 4**

| Oligo | Sequence (5′ – 3′) |
|---|---|
| **ND-NCP oligos** | **EMSAs** |
| ND_001 | ATCGGATGTATATATCTGACACGTGCCTGGAGACTAGGGAGTAATCCCCTTGGCGG<br>TTAAAACGCGGGGGACAGCGCGTACGTGCGTTTAAGCGGTGCTAGAGCTGTCTACG<br>ACCAATTGAGCGGCCTCGGCACCGGGATTCTCGAT |
| ND_002 | /56-FAM/ ATCGAGAATCCCGGTGCCGAGGCCGCTCAATTGGTCGTAGACAGC<br>TCTAGCACCGCTTAAACGCACGTACGCGCTGTCCCCCGCGTTTTAACCGCCAAGGG<br>GATTACTCCCTAGTCTCCAGGCACGTGTCAGATATATACATCCGAT |
| **Linear DNA oligos** | **STK** |
| Linear DNA_001 | ATCGGATGTATATATCTGACACGTGCCTGGAGACTAGGGAGTAATCCCCTTGGCGG<br>TTAAAACGCGGGGGACAGCGCGTACGTGCGTTTAAGCGGTGCTAGAGCTGTCTACG<br>ACCAATTGAGCGGCCTCGGCACCGGGATTCTCGAT |
| Linear DNA_002 | /56-FAM/ATCGAGAATCCCGGTGCCGAGGCCGCTCAATTGGTCGTAGACAGC<br>TCTAGCACCGCTTAAACGCACGTACGCGCTGTCCCCCGCGTTTTAACCGCCAAGGG<br>GATTACTCCCTAGTCT |
| Linear DNA_003 | /5Phos/CAGGCACGTGTCAGATATATACATCCGAT |
| **Gap-NCP$_{-5.5}$ oligos** | **Cryo-EM (without /56-FAM/), STK, and EMSAs** |
| Gap-NCP$_{-5.5}$_001 | ATCGGATGTATATATCTGACACGTGCCTGGAGACTAGGGAGTAATCCCCTTGGCGG<br>TTAAAACGCGGGGGACAGCGCGTACGTGCGTTTAAGCGGTGCTAGAGCTGTCTACG<br>ACCAATTGAGCGGCCTCGGCACCGGGATTCTCGAT |
| Gap-NCP$_{-5.5}$_002* | /56-FAM/ATCGAGAATCCCGGTGCCGAGGCCGCTCAATTGGTCGTAGACAGC<br>TCTAGCACCGCTTAAACGCACGTACGCGCTGTCCCCCGCGTTTTAACCGCCAAGGG<br>GATTACTCCCTAGTCTCCAGGCACGT |
| Gap-NCP$_{-5.5}$_003 | /5Phos/TCAGATATATCCATCCGAT |
| **Gap-NCP$_{-4.5}$ oligos** | **Cryo-EM (without /56-FAM/), STK, and EMSAs** |
| Gap-NCP$_{-4.5}$_001 | ATCGGATGTATATATCTGACACGTGCCTGGAGACTAGGGAGTAATCCCCTTGGCGG<br>TTAAAACGCGGGGGACAGCGCGTACGTGCGTTTAAGCGGTGCTAGAGCTGTCTACG<br>ACCAATTGAGCGGCCTCGGCACCGGGATTCTCGAT |
| Gap-NCP$_{-4.5}$_002* | /56-FAM/ATCGAGAATCCCGGTGCCGAGGCCGCTCAATTGGTCGTAGACAGC<br>TCTAGCACCGCTTAAACGCACGTACGCGCTGTCCCCCGCGTTTTAACCGCCAAGGG<br>GATTACTCCCTAGTCT |
| Gap-NCP$_{-4.5}$_003 | /5Phos/CAGGCACGTGTCAGATATATACATCCGAT |
| **Gap-NCP$_{-3.5}$ oligos** | **Cryo-EM (without /56-FAM/), STK, and EMSAs** |
| Gap-NCP$_{-3.5}$_001 | ATCGGATGTATATATCTGACACGTGCCTGGAGACTAGGGAGTAATCCCCTTGGCGG<br>TTAAAACGCGGGGGACAGCGCGTACGTGCGTTTAAGCGGTGCTAGAGCTGTCTACG<br>ACCAATTGAGCGGCCTCGGCACCGGGATTCTCGAT |
| Gap-NCP$_{-3.5}$_002* | /56-FAM/ATCGAGAATCCCGGTGCCGAGGCCGCTCAATTGGTCGTAGACAGC<br>TCTAGCACCGCTTAAACGCACGTACGCGCTGTCCCCCGCGTTTTAACCGCCAAGGG<br>GATTA |
| Gap-NCP$_{-3.5}$_003 | /5Phos/TCCCTAGTCTCCAGGCACGTGTCAGATATATCCATCCGAT |
| **Gap-NCP$_{-2.5}$ oligos** | **STK and EMSAs** |
| Gap-NCP$_{-2.5}$_001 | ATCGGATGTATATATCTGACACGTGCCTGGAGACTAGGGAGTAATCCCCTTGGCGG<br>TTAAAACGCGGGGGACAGCGCGTACGTGCGTTTAAGCGGTGCTAGAGCTGTCTACG<br>ACCAATTGAGCGGCCTCGGCACCGGGATTCTCGAT |
| Gap-NCP$_{-2.5}$_002* | /56-FAM/ATCGAGAATCCCGGTGCCGAGGCCGCTCAATTGGTCGTAGACAGC<br>TCTAGCACCGCTTAAACGCACGTACGCGCTGTCCCCCGCGTTTTAACCGC |
| Gap-NCP$_{-2.5}$_003 | /5Phos/AAGGGGATTACTCCCTAGTCTCCAGGCACGTGTCAGATATATCCATCCGAT |
| **Gap-NCP$_{-1.5}$ oligos** | **STK and EMSAs** |

| | |
|---|---|
| Gap-NCP$_{-1.5}$_001 | ATCGGATGTATATATCTGACACGTGCCTGGAGACTAGGGAGTAATCCCCTTGGCGGTTAAAACGCGGGGGACAGCGCGTACGTGCGTTTAAGCGGTGCTAGAGCTGTCTACGACCAATTGAGCGGCCTCGGCACCGGGATTCTCGAT |
| Gap-NCP$_{-1.5}$_002* | /56-FAM/ATCGAGAATCCCGGTGCCGAGGCCGCTCAATTGGTCGTAGACAGCTCTAGCACCGCTTAAACGCACGTACGCGCTGTCCCCCGCGT |
| Gap-NCP$_{-1.5}$_003 | /5Phos/TTAACCGCCAAGGGGATTACTCCCTAGTCTCCAGGCACGTGTCAGATATATCCATCCGAT |

| **Gap-NCP$_{-4.5+2}$ oligos** | **STK and EMSAs** |
|---|---|
| Gap-NCP$_{-4.5+2}$_001 | ATCGGATGTATATATCTGACACGTGCCTGGAGACTAGGGAGTAATCCCCTTGGCGGTTAAAACGCGGGGGACAGCGCGTACGTGCGTTTAAGCGGTGCTAGAGCTGTCTACGACCAATTGAGCGGCCTCGGCACCGGGATTCTCGAT |
| Gap-NCP$_{-4.5+2}$_002* | /56-FAM/ATCGAGAATCCCGGTGCCGAGGCCGCTCAATTGGTCGTAGACAGCTCTAGCACCGCTTAAACGCACGTACGCGCTGTCCCCCGCGTTTTAACCGCCAAGGGGATTACTCCCTAGT |
| Gap-NCP$_{-4.5+2}$_003 | /5Phos/ TCCAGGCACGTGTCAGATATATCCATCCGAT |

| **Gap-NCP$_{-4.5+3}$ oligos** | **STK and EMSAs** |
|---|---|
| Gap-NCP$_{-4.5+3}$_001 | ATCGGATGTATATATCTGACACGTGCCTGGAGACTAGGGAGTAATCCCCTTGGCGGTTAAAACGCGGGGGACAGCGCGTACGTGCGTTTAAGCGGTGCTAGAGCTGTCTACGACCAATTGAGCGGCCTCGGCACCGGGATTCTCGAT |
| Gap-NCP$_{-4.5+3}$_002* | /56-FAM/ATCGAGAATCCCGGTGCCGAGGCCGCTCAATTGGTCGTAGACAGCTCTAGCACCGCTTAAACGCACGTACGCGCTGTCCCCCGCGTTTTAACCGCCAAGGGGATTACTCCCTAG |
| Gap-NCP$_{-4.5+3}$_003 | /5Phos/CTCCAGGCACGTGTCAGATATATCCATCCGAT |

| **Gap-NCP$_{-4.5+4}$ oligos** | **STK and EMSAs** |
|---|---|
| Gap-NCP$_{-4.5+4}$ _001 | ATCGGATGTATATATCTGACACGTGCCTGGAGACTAGGGAGTAATCCCCTTGGCGGTTAAAACGCGGGGGACAGCGCGTACGTGCGTTTAAGCGGTGCTAGAGCTGTCTACGACCAATTGAGCGGCCTCGGCACCGGGATTCTCGAT |
| Gap-NCP$_{-4.5+4}$ _002* | /56-FAM/ATCGAGAATCCCGGTGCCGAGGCCGCTCAATTGGTCGTAGACAGCTCTAGCACCGCTTAAACGCACGTACGCGCTGTCCCCCGCGTTTTAACCGCCAAGGGGATTACTCCCTA |
| Gap-NCP$_{-4.5+4}$ _003 | /5Phos/TCTCCAGGCACGTGTCAGATATATCCATCCGAT |
